# Competitive exclusion of uropathogenic *E. coli* biofilm by *Salmonella* through matrix inhibition

**DOI:** 10.1101/2020.01.18.911263

**Authors:** Sandeep Miryala, S Chandramohan, C. S. Srinandan

## Abstract

Biofilm is a predominant lifestyle of bacteria in host and non-host environments with cell collectives and extracellular matrix as the defining principles of biofilm. Several factors trigger biofilm formation including response to competition. Urinary tract infections (UTI) are highly prevalent worldwide and mainly caused by uropathogenic *E. coli* (UPEC), which progresses into chronic form due to the biofilm formation by the pathogen. In this study, we hypothesized that competition for territorial space could occur between species by intervening in the biofilm matrix production, particularly of UPEC, thereby reducing its colonizing ability. UPEC colony displays different morphology in congo red media based on matrix production, which we exploited for screening bacterial isolates capable of inhibiting the matrix. This was validated by using the cell-free supernatants of the isolates to impair UPEC biofilm. Isolates that inhibited matrix production belonged to species of Shigella, Escherichia, Enterobacter, and Salmonella from Enterobacteriaceae family. Competition experiments between the isolates and UPEC revealed spiteful interactions particularly during biofilm formation, indicating fierce competition for territorial space colonization. The isolate *Salmonella enterica* B1 could competitively exclude UPEC in the biofilm. Altogether, we show that interference competition by matrix inhibition occurs as a strategy by bacteria to colonize territorial space.

## Introduction

Urinary tract infections (UTIs) are prevalent in large scale among the human population, with about 150 million people worldwide getting infected by UTI annually (Flores-Mireles *et al.,* 2015). Uropathogenic *E. coli* (UPEC) is the predominant causative agent in UTI and recurrent UTI (rUTI) is a common and challenging problem causing substantial morbidity (Glover *et al.,* 2014). Biofilm plays a key role in UPEC pathogenesis that cause persistence of infection (Soto *et al*., 2006; Tamadonfar *et al*., 2019). Biofilm formation on urinary catheters are a significant problem globally that is responsible for 40% of nosocomial infections and is extremely difficult to treat (Walker *et al*., 2019).

Biofilm matrix acts as a physical barrier to protect the cells from predation, radiation, desiccation, resistance/tolerance towards the antimicrobials including cells of the immune system, and matrix also provide biofilm cells the advantage in accessing nutrients and other communal benefits (Xavier and Foster, 2007; Leid 2009; DePas *et al*., 2014; Srinandan *et al*., 2015; Dragoš, and Kovács, 2017). The important matrix components of UPEC are curli and cellulose, where curli are amyloid proteins that helps in adhesion, cell-surface interaction cell-cell interactions and acts as structural scaffold to promote biofilm assembly (Shanmugam *et al*., 2019). On the other hand, cellulose provides the elastic behavior, 3D structure, tolerance to chlorine, and spatial assortment in the biofilm (Solano et al., 2002; Srinandan *et al*., 2015; Serra and Hengge, 2019). A simple *in vitro* assay exists to score the production of curli and cellulose in *E. coli* colonies wherein, the congo red dye is added to stain the matrix components (Serra and Hengge, 2017).

The life of *E. coli* populations is biphasic, that is, it must adapt and survive both in host and non-host environments. UPEC is found in wastewaters even after treatment that is let off into natural water bodies or soil (Anastasi *et al*., 2012; Zhi *et al*., 2019). However, the persistence of UPEC in nature is not very clear, though *E. coli* populations establish in the soil or water environments (Blount 2015). In non-host environmental conditions which is stressful and fluctuating, biofilm is the plausible lifestyle of UPEC survival (DePas *et al*., 2014). Around 40%-80% of bacteria survive as biofilms in nature, making it the predominant lifestyle (Flemming and Wuertz, 2019). Sociobiological interactions are rich in the spatially structured biofilm, among which competition between species occurs for finite resources. However, Oliveira *et al*., (2015) showed that biofilm formation itself is a strategic lifestyle of the cells in response to competition. Thus, it’s imperative that microbial species would compete for territorial colonization by forming biofilm, and as matrix is important for biofilm formation, we hypothesized that one species may secrete compounds to inhibit matrix production of the competing species. If there is such kind of competition, the species that inhibit matrix production of UPEC could potentially be used in biotechnological applications to control UPEC. Therefore, in this study, we attempted to screen matrix-inhibiting bacteria against UPEC by using the Congo red method and with further testing, we gain some insights on competitive exclusion of UPEC by Enterobacteriaceae family.

## Results and Discussion

Matrix is important for biofilm lifestyle (Flemming *et al*., 2016), devoid of which bacterial cell collectives lose the critical features of biofilm like resilience to stress, social interactions, architecture, etc. In this study, we attempted to isolate matrix-inhibiting bacteria of UPEC from the soil samples.

### Inhibition of UPEC biofilm matrix production by soil bacterial isolates

We designed a novel and simple methodology to screen for bacterial isolates that could specifically suppress the matrix production in UPEC thereby inhibiting biofilm formation for which we used a traditional congo red (CR) dye-containing media. The predominant matrix components in *E. coli* and *Salmonella* are cellulose and curli proteins (Serra and Hengge, 2017). The CR dye binds to these matrix components to give red color to the colony and when there is no matrix, it displays a white colony. The former develops a rough colony due to the matrix components and the latter forms a smooth colony due the absence of cellulose and curli (Serra and Hengge, 2017). Thirty different soil samples near wastewaters were sampled in a sterile container, serially diluted in PBS and, plated on CR agar. The UPEC was spotted and we observed color of the colony after incubation for three days **(Figure 1a)**. The UPEC colonies that showed smooth and white (SAW) morphology were further selected and the peripheral bacterial cells from these colonies were collected, purified by traditional streak-plate method, and validated with the CR plate assay. Seven bacterial isolates showed positive results by apparently inhibiting the matrix-production in UPEC, which were named as A1, B1, C1, F1, P1, T1 and Z1 **(Figure 1b)**.

**Figure 1.**
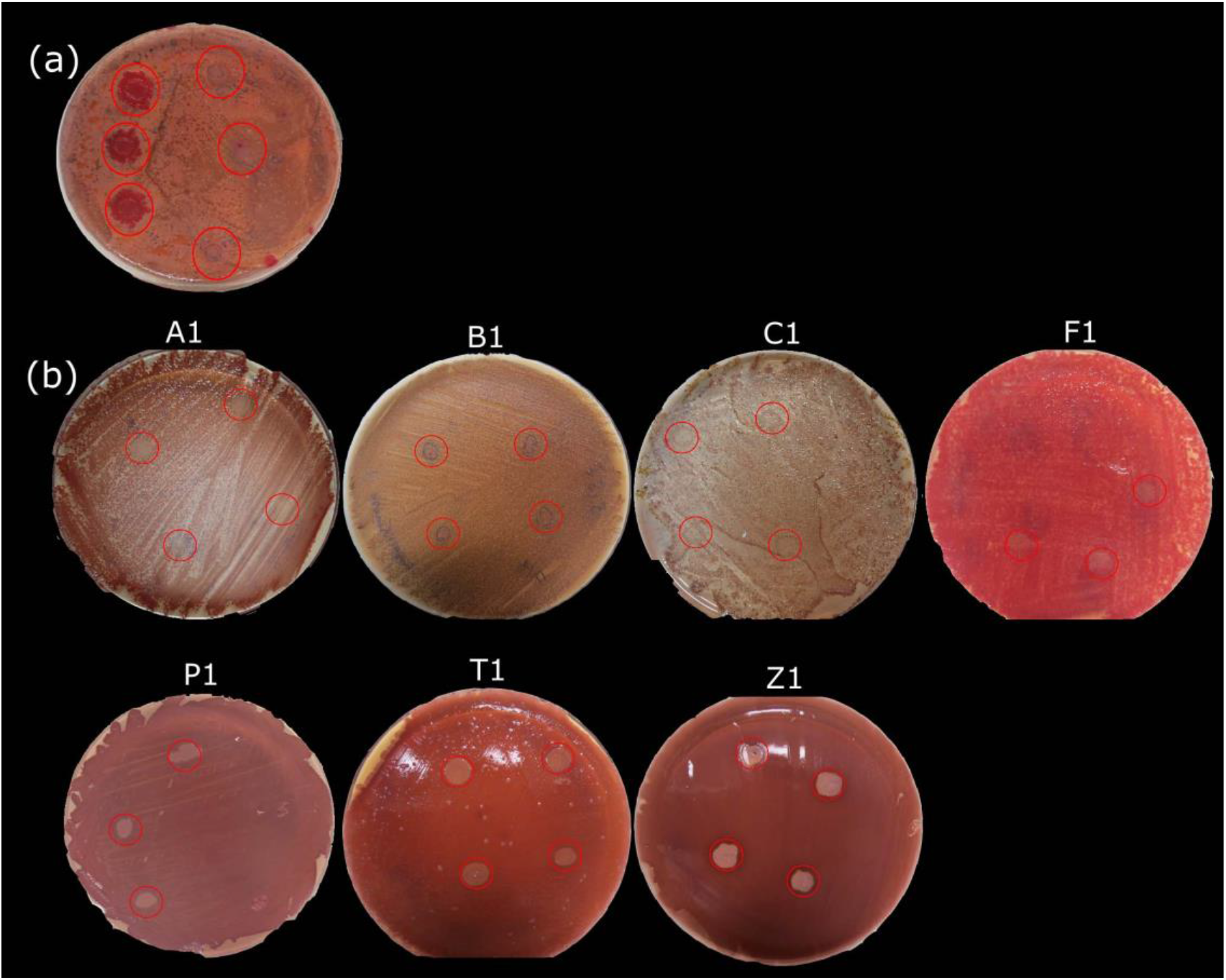
Screening for bacterial isolates on congo red (CR) media that could potentially inhibit matrix production in UPEC biofilms. Representative plates showing the (a) influence of soil bacterial isolates and (b) influence of pure culture of bacterial isolates. Alphabets on the plates refer to the name of the isolates. UPEC colonies in the CR plate are marked by red circle.

### Culture supernatant of the isolates act against UPEC biofilm

Further, we asked if the interference in UPEC matrix production could be due to the competition sensing (Cornforth and Foster, 2013), for which we collected the cell-free supernatant from colony (CFSC) of the isolates that were grown in the proximity of UPEC colony. The planktonic growth of UPEC was affected by the CFSC of the isolates (Figure S1). Except for the CFSC of isolate A1, all other extracts had a highly significant inhibitory effect on the UPEC planktonic growth (Mann-Whitney *U* test, *P* < 0.001, *n* = 5). However, the CFSC of all the isolates displayed a significant inhibitory effect on biofilm formation of UPEC (Mann-Whitney *U* test, *P* < 0.001, *n* = 5), indicating that inhibition of matrix production has a direct consequence on biofilm formation (Figure S1).

We also tested if the independently grown culture supernatant of the isolates without competition sensing (henceforth called Competition-Sensing Independent Supernatant or CSIS) has effect on growth and cell assemblages of UPEC. The CSIS were tested for its effect on UPEC biofilm formation at three different concentration, whereby all the three concentrations substantially inhibited biofilm formation (Figure S2), but we used 25% for further studies. The CSIS of the isolate F1 had inhibited the planktonic growth by more than 60%, however other culture supernatants of the bacterial isolates inhibited 20%-50% of the planktonic growth of UPEC (Figure 2a). Less than 50% of adhesion of UPEC was inhibited by the CSIS (Figure 2b). But, more than 90% of biofilm formation was prevented and more than 70% of biofilm eradication was seen with the CSIS of all the isolates (Figure 2c and d). Absolute values corresponding to Figures a-d are shown in Figure S3. The formation of biofilm and dispersal of preformed biofilm, particularly the submerged biofilm on glass surfaces were also effectively inhibited or eradicated respectively by the CSIS from all the isolates as visualized by fluorescence microscopy (Figure 2e and S4).

**Figure 2.**
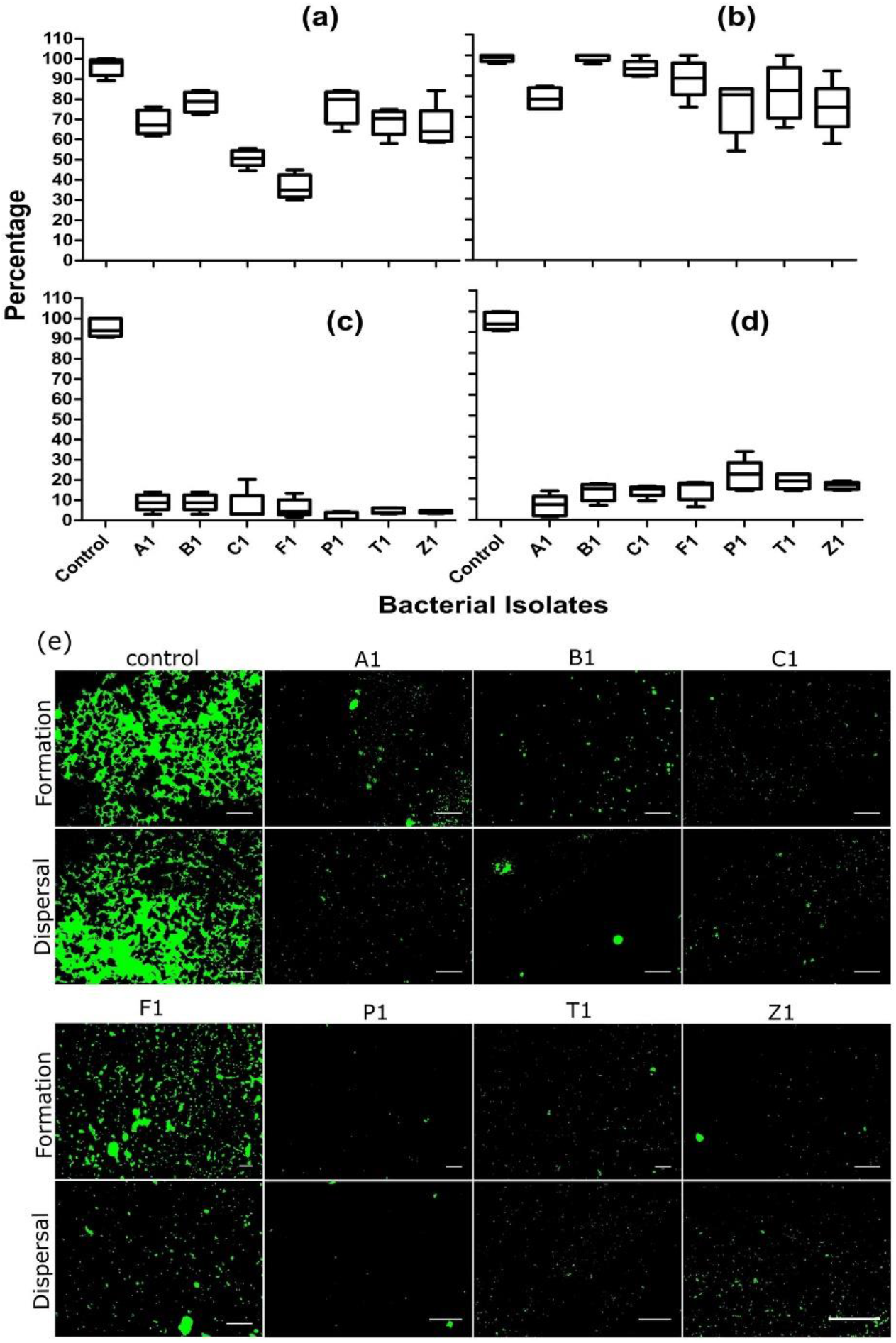
Influence of bacterial isolates on partially submerged biofilm (PSM) (a) planktonic growth, (b) adhesion, (c) biofilm formation and, (d) dispersal of UPEC biofilm. *n* = 5. The absolute values of absorbance are given in Figure S3. (e) Representative images of fluorescent microscopy showing biofilm formation and dispersal of preformed biofilm. Scale bar = 250 μm. Quantified data of images are shown in Figure S4.

Biofilm matrix has many functions among which providing the structural support to the cell assemblages is one of the most important. Inhibition of the biofilm matrix production may be involved in inhibiting the UPEC biofilm, but we also observed an effective eradication of preformed biofilm. Interference competition takes place between microbial species leading to a chemical warfare between them during competition (Ghoul and Mitri, 2016). The warfare-related chemical compounds are released out of the producing cell to kill or inhibit the competing species. Thus, cell-free culture supernatants of bacteria are a rich source of bioactive compounds that could be exploited in biotechnology. Antibiofilm compounds are also being discovered using the culture supernatants (Valle *et al*., 2006; Nithya and Pandian 2010). We observed that the matrix-inhibiting compounds by the selected isolates were produced even without sensing the competition (Figure 2), indicating that these compounds may have other roles too. Competition sensing hypothesis proposed by Cornforth and Foster (2013) predicts that the physiological response evolves due to ecological competition. However, in this case the matrix inhibition of UPEC was not in response, but it could be a physiologically produced metabolite having multiple roles, similar to the phenazines (Whelan *et al*., 2006).

### Physicochemical nature of the culture supernatant

To find the preliminary physicochemical nature of CSIS, it was treated with 2-β mercaptoethanol (BME), trypsin, proteinase K and heat. The treated supernatants were used to check their effect on matrix production in CR plates, planktonic growth, and biofilm formation of UPEC. Many isolates lost the capacity to inhibit both cellulose and curli production after domestication in the laboratory media. However, the culture supernatants of the isolates, A1, P1, T1 and, Z1 turned the proximate colony of UPEC into pink color (Figure S5a), which indicates that only cellulose is expressed but not curli (Serra and Hengge, 2017). The culture supernatants from the isolate B1 was consistent in suppressing both cellulose and curli of UPEC (Figure S5a). The culture supernatants of A1, B1, P1 and, Z1 when treated with heat, lost its capacity to influence the matrix production of UPEC (Figure S5a). However, in some instances, the treated culture supernatant had inhibited 70%-90% of planktonic growth particularly after subjecting it to heat from B1, T1 and Z1, which could also be seen as zone of inhibition in B1 and T1 in the CR plates (Figure S5a and b). Trypsin treatment rescued nearly 40% of the inhibitory effect of culture supernatant from isolate B1, but matrix production was marginally less than control (Figure S5a and c). BME treatment of culture supernatant of isolate C1 rescued biofilm inhibition, which was also observed in the rescuing matrix production (Figure S5a and c). The extract of isolate F1 showed a zone of inhibition indicating growth inhibition that was consistent with the planktonic growth inhibitory activity (Figure S5a and b). Treatment of the extract subjected with BME from isolate P1 also inhibited more than 70% of planktonic growth of UPEC (Figure S5b). However, more than 50% of matrix production was rescued when the culture supernatant of P1 was heat-treated. Altogether, these results showed that matrix suppression and biofilm inhibition from the culture supernatants is not by proteins except for the supernatant of isolate C1 (Figure S5). We speculate that the mechanism of inhibition of UPEC matrix or biofilm could be by producing specific polysaccharides that inhibit matrix gene regulation, similar to that reported Valle et al., (2006) or some small molecules, which are sensitive to heat. Interference of these molecules in c-di-GMP signaling cannot be ruled out, as the higher intracellular concentrations of c-di-GMP activate matrix production (Qvortrup *et al*., 2019).

### The bacterial isolates that inhibit matrix production belong to Enterobacteriaceae

The seven bacterial isolates that showed matrix-inhibiting activity of UPEC were identified by using 16S rRNA gene sequencing and analyzing its phylogeny (Figure 3). The 16S rRNA sequences were submitted to NCBI with accession numbers as shown in Table S1. The isolates were submitted to National Centre for Microbial Resource, Pune, India (Table S1). The isolate A1 was identified as *Escherichia fergusonii,* B1 as *Salmonella enterica,* C1 belonged to the genus *Escherichia,* F1 was *E. fergusonii,* isolates P1 and T1 were *Shigella flexneri,* and the isolate Z1 was identified as *Enterobacter cloacae.* All these isolates belonged to Enterobacteriaceae family. Two general kinds of competition occurs between species, (a) exploitative competition, where the resources could be highly exploited by one species thus reducing the fitness payoff in the other and, (b) interference competition, where one species interferes directly into the growth of other species by inhibiting or killing it (Ghoul and Mitri, 2016). Ecological competition among the Enterobacteriaceae family is intense because the resources used by its members are similar. For example, exploitative competition for iron emerges between Enterobacteriaceae group, in which the species compete by the production of siderophores (Deriu *et al*., 2014). Litwak *et al*., (2019) shows that they compete each other for oxygen in the gut environment. Also, fierce interference competition occurs among different species of this group by producing colicins and microcins (Nedialkova *et al*., 2014; Sassone-Corsi *et al*., 2016).

**Figure 3.**
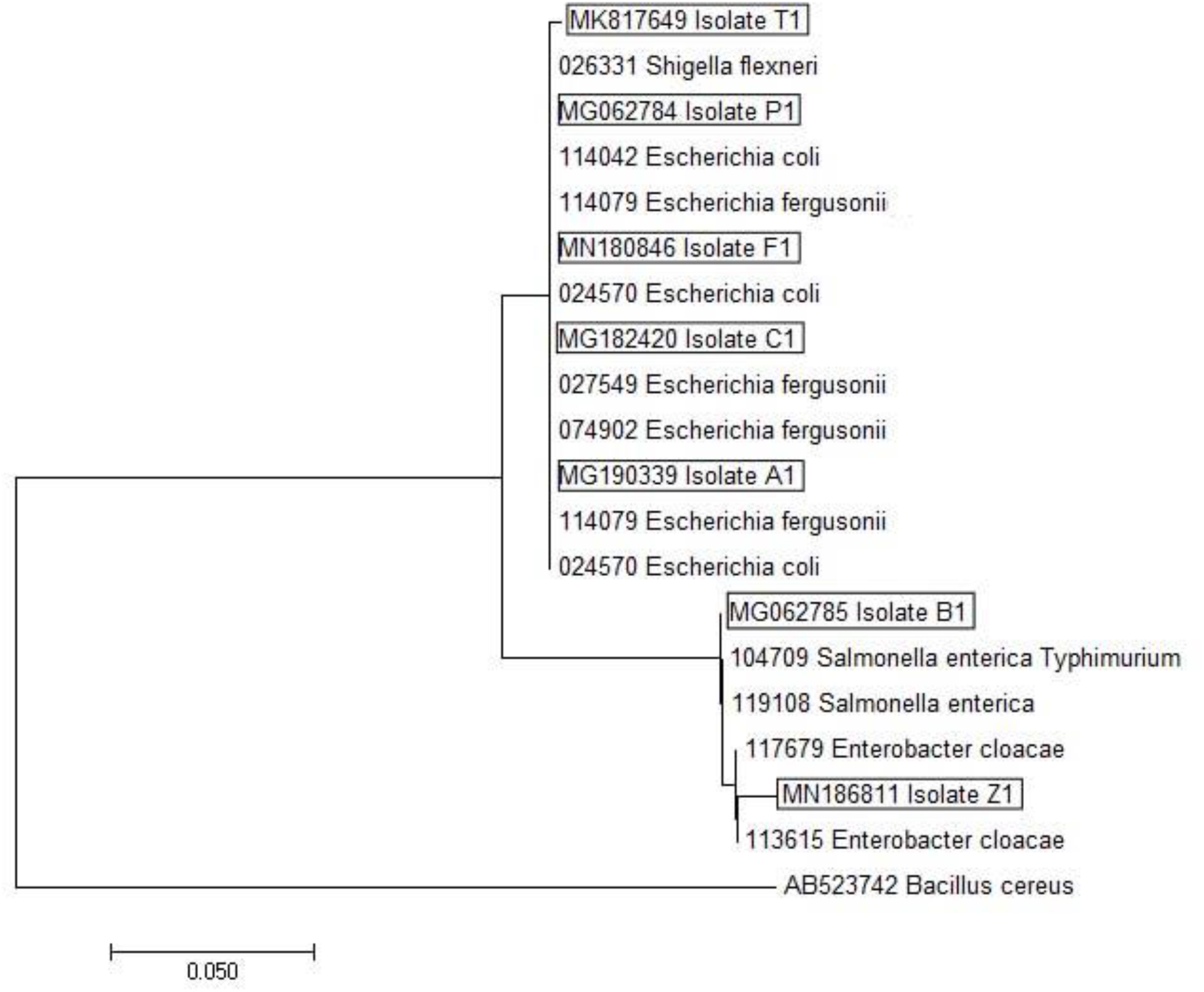
Phylogenetic tree of the isolates that show biofilm matrix inhibiting activity against UPEC. NCBI accession number is depicted before the isolate or organism name.

### Territorial conflict results in tragedy of commons

Further, competition experiments were performed with UPEC against all the isolates. The relative fitness between the monoculture and coculture of the organism was calculated, as it gives an insight into the kind of interaction in the coculture between the UPEC and the isolate. A relative fitness value ≈1 indicates neutral interaction between both, and a value <1 for both species indicate cooperation, due to increased fitness payoff in coculture than monoculture. If either of the organism in the coculture has a relative fitness value <1, then it is possibly exploiting the other for its benefit. If the relative fitness value of both the isolate and UPEC is >1, it indicates tragedy of commons, where both the organisms reduced its absolute fitness when in coculture.

In planktonic growth, the relative fitness of both UPEC and the isolate was significantly >1 in the case of A1 and T1, suggesting both the organisms tragedized during coculture (Figure 4a, S6a and S6e), although the isolate T1 was relatively fitter than UPEC (Figure 4c). The fitness of UPEC monoculture was significantly higher than coculture, with B1 and C1 isolates (Figure 4a, S6b and S6c), but the fitness of isolates in either mono or coculture did not change. However, the fitness of B1 and C1 were significantly higher in the coculture than UPEC. Isolate P1 and UPEC possibly had a neutral interaction, thus no significant positive or negative payoff was observed on its fitness (Figure 4a and S6d). Coculture of UPEC with the isolate Z1 enhanced the fitness payoff of UPEC than monoculture but reduced the payoff of Z1 in coculture relatively to the monoculture (*n*=4, *P*<0.01, one sample *t* test) (Figure 4a). Due to growth inconsistencies, competition experiments with isolate F1 was not determined.

**Figure 4.**
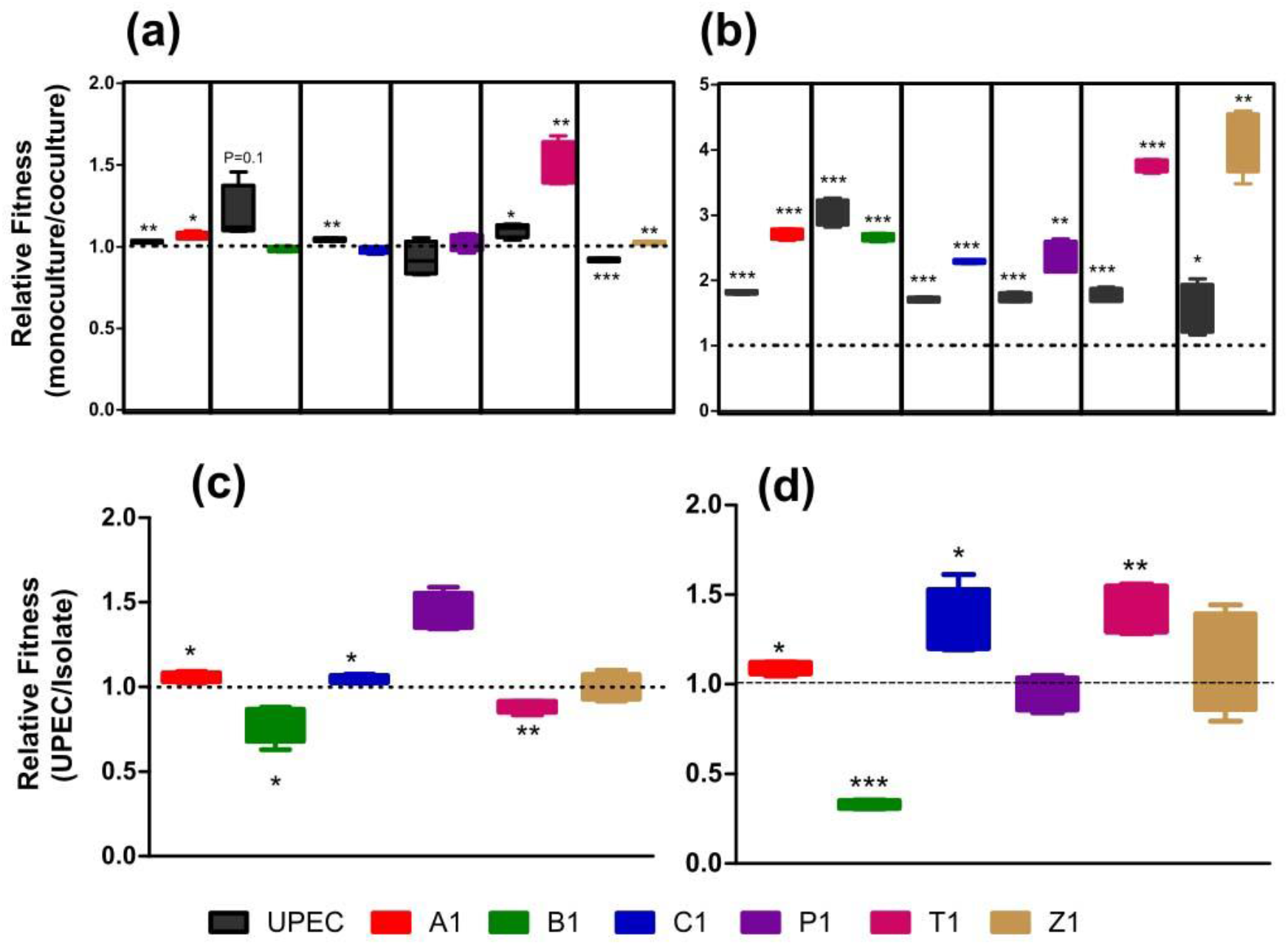
Relative fitness of UPEC and the bacterial isolate in monoculture and coculture with respective isolate during (a) planktonic growth and (b) biofilm growth. Relative fitness of the organisms in coculture experiments during (c) planktonic growth and (d) biofilm growth. The color codes suggest that the coculture experiments were performed with that isolate and the UPEC. *n* = 4, one sample *t* test was performed to determine significance with a theoretical mean of 1.00 (\**P*<0.05, \*\**P*<0.01, \*\*\**P*<0.001). Absolute fitness values are shown in Figure S6.

In biofilm growth, fitness payoff significantly reduced for all the cultures in coculture biofilm relative to the monoculture (Figure 4b and S6). This clearly shows that a fierce competition occurs between the isolates and UPEC. As all the cultures belong to the Enterobacteriaceae family, which share similar resources, spiteful competition possibly occurred for the territorial space. However, relative fitness is higher for UPEC when there is competition between the isolates, A1, C1 and, T1 (Figure 4d). The relative fitness in competition with P1 and Z1 was nearly 1.0 (Figure 4d). Nevertheless, the isolate B1 emerged with higher fitness value relatively than UPEC even in biofilm growth (Figure 4d).

Matrix production by the species push the biofilm cells upwards, which can access more oxygen and nutrients and suffocate the non-producers (Xavier and Foster, 2007). Thus, matrix producers will have positive fitness payoff than the non-producers resulting in colonization of territory by producers. Significant reduction of fitness payoff in coculture than monoculture between the isolates and UPEC for territorial colonization of the surface indicate tragedy of commons. The plastic surface of the microtiter plate’s well is the intact common good and, matrix secretion favors the producer to colonize the surface.

### Salmonella enterica B1 competitively excludes UPEC in biofilm

As the isolate B1, which was identified as *Salmonella enterica* (Figure 3) exhibited matrix inhibition and higher fitness in competition experiments with UPEC (Figure 4), we transformed plasmids expressing fluorescent proteins in both *S. enterica* B1 and the UPEC to observe microscopically the spatial arrangement of the cell types. *S. enterica* B1 outcompeted UPEC in the submerged biofilm on glass slide (Figure 5a). Quantification of the images revealed that the biomass and substratum coverage was higher for UPEC in the monoculture biofilm that significantly reduced in coculture (Figure 5b and S7). Biomass and substratum coverage increased significantly to *S. enterica* B1 in the coculture than monoculture (Figure 5b and S7). In submerged biofilm, the sum total of biomass was significantly high in coculture than the sum of monoculture of both organisms (Mann-Whitney *U* test, *P* < 0.02, *n* = >20) (Figure S7), implying that the *S. enterica* B1 and UPEC could increase their overall productivity in submerged biofilm, but *S. enterica* B1 predominates. Monoculture productivity of the *S. enterica* B1 was significantly lesser than UPEC, but *S. enterica* B1 increased its biomass in coculture (Figure S7a). The *S. enterica* B1 also significantly increased its substratum coverage in coculture than the monoculture (Mann-Whitney *U* test, *P* < 0.02, *n* = >20), though the overall coverage of both mono and coculture was similar (Figure S7b). We speculate that the matrix inhibition of UPEC might have favored the *S. enterica* B1 to colonize the surface, thus suffocating UPEC in the biofilm, similar to the model proposed by Xavier and Foster (2007). Conflicts between Salmonella and *E. coli* in different contexts have been reported (Nedialkova *et al*., 2014; Sassone-Corsi *et al*., 2016; Deriu *et al*., 2014; Litwak *et al*., 2019), but here we observed the territorial conflict among these two species in the context of biofilm formation, where the *S. enterica* B1 competitively excluded UPEC.

**Figure 5.**
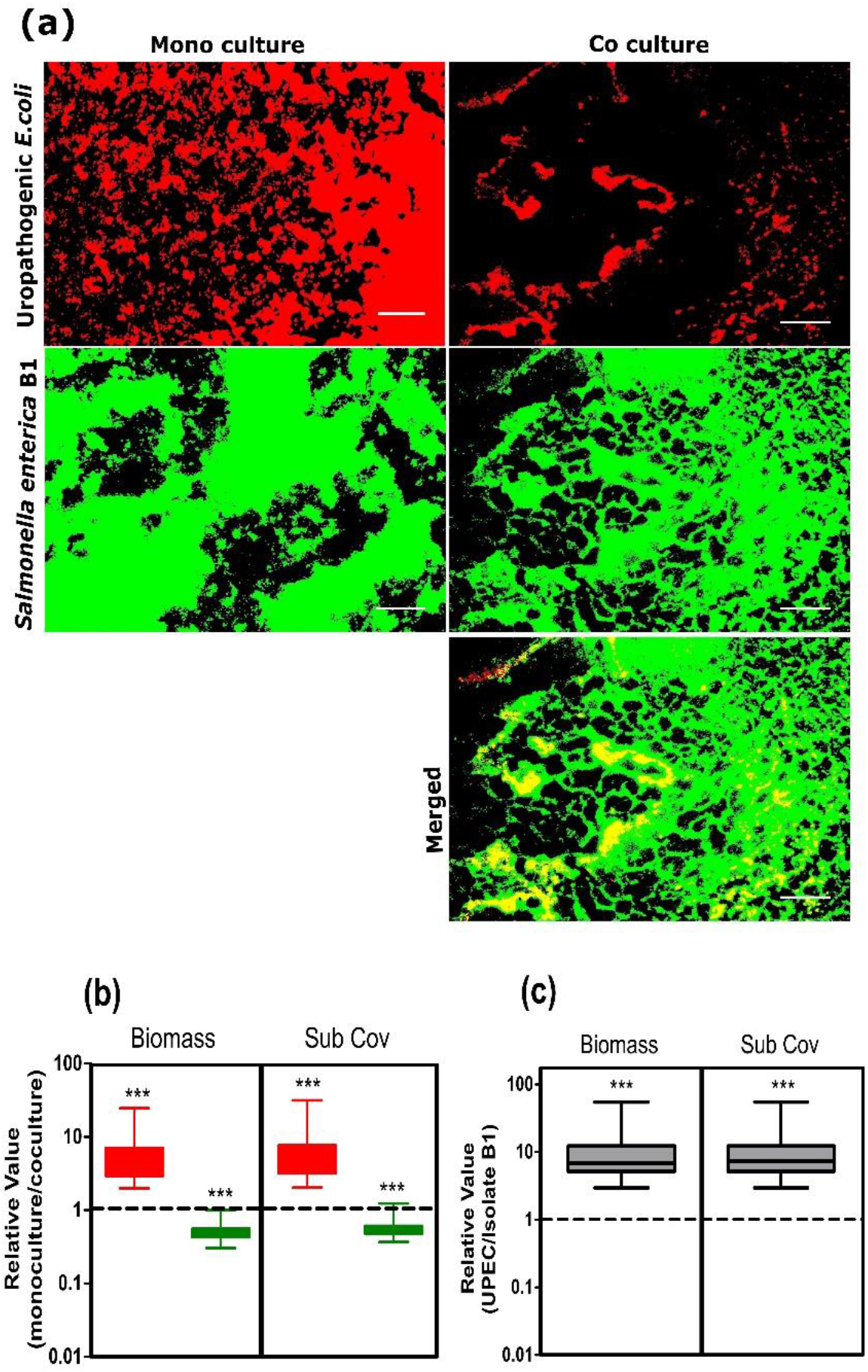
Competition in biofilm between UPEC and the *S. enterica* B1 (a) Representative fluorescence microscopic biofilm images in monoculture and coculture. Scale bar is 250 μm. Quantified data of the fluorescence microscopic biofilm images showing relative values of biomass and substratum coverage (sub cov) with respect to (b) monoculture is to coculture and (c) UPEC is to *S. enterica* B1 in the coculture. Red color in the image and graph is represented as UPEC and green is represented with the *S. enterica* B1. *n* = >20. One sample *t* test with a hypothetical value of 1.0 was performed to determine the significance (*** *P*<0.001).

## Concluding Remarks

The predominant survival strategy of bacteria in host and non-host conditions are as biofilm. Matrix is the most important component for biofilm bacteria to colonize surfaces, thus targeting it will be a superior strategy to treat pathogenic biofilms. However, natural products released by microbes during competition for territorial space could be potentially exploited for discovering novel antibiofilm compounds. Modern idea of infection therapy is based on antibiotics, which was discovered by Alexander Fleming and currently, around 69% of antimicrobials are from natural products (Pham *et al*., 2019). Also, there is a renewed interest in discovering drugs from natural products (Waldetoft *et al*., 2019). The simple screening assay that we developed in this study to isolate bacteria that could inhibit the matrix production in competition for territorial space based on the congo red agar test, could potentially be used for high-throughput screening of natural antibiofilm compounds (Figure 1).

Our study also gives an insight that bacterial species may compete for territorial colonization by inhibiting matrix, thus suppressing biofilm of the competing species. Particularly, among the species that ecologically compete for similar resources, in this case the Enterobacteriaceae family. The common resource for the species is the substratum surface and competitive exclusion of UPEC was observed by some members of Enterobacteriaceae where Salmonella was more effective. In this era of antibiotic resistance, such strategies, where non-pathogenic species that competitively exclude pathogen colonization by intervening in its matrix production could potentially be screened to develop probiotics.

## Experimental Procedures

### Bacterial strain and growth conditions

Uropathogenic *E. coli* UTI89 (henceforth referred to as UPEC) strain (gifted by Prof. Matthew A. Mulvey, University of Utah) was used for all the experiments. The media used for culturing was Yeast Extract Casamino Acids (YESCA) (Yeast Extract 0.5g/L Casamino Acids 10g/L) (Wu et al. 2012).

### Screening for bacterial strains that could inhibit biofilm matrix of UPEC

In YESCA media supplemented with Congo red dye, 40 μg mL^-1^ and Coomassie brilliant blue, 20 μg mL^-1^, biofilm matrix producing *E. coli* forms red, dry and rough morphology (RDAR) and absence of matrix production will give smooth and white color (SAW) (Serra and Hengge, 2017). This was used as an assay for screening bacteria that could potentially inhibit matrix production in UPEC. Several soil samples near domestic wastewater in Thanjavur, Tamil Nadu, India were collected in a sterile container, which were serially diluted in sterile Phosphate Buffered Saline (PBS) and spread plated on congo red (CR) agar plate. Around 10 μl of PBS-washed overnight-grown UPEC culture was spotted on the CR agar plate that was incubated for 3 days at 25 °C. Surrounding colonies of the UPEC that formed SAW morphotype were picked and isolated in LB medium. Pure colonies of bacterial isolates were validated for their influence on UPEC colony morphology by the same method.

### Preparation and physicochemical analysis of culture supernatants

The supernatant from pure culture of selected soil isolates were prepared according to Farmer *et al*., (2014) but with modifications as follows. We used a cell-free supernatant from colony (CFSC) or Competition-Sensing Independent Supernatant (CSIS) wherein, the CFSC was collected by growing a lawn of the bacterial isolate in YESCA agar, spotting UPEC in the plate and, incubating it for three days. The bacterial isolate’s cells that were in proximity to the UPEC spot were scraped with the pipette tip and added in sterile YESCA broth, which was centrifuged at 5000 rpm for 10 mins and filter sterilized with a 0.22 μm nylon-66 membrane (HiMedia). For CSIS, the bacterial isolate’s cells were grown in YESCA broth for 3 days at 25°C in static condition, which was centrifuged at 5000 rpm for 10 mins and filter sterilized (HiMedia). The resulting culture supernatants were stored at 4°C and used for further experiments.

The cell-free supernatants were treated with either 2-Mercaptoethanol (BME) (SRL), Trypsin, Proteinase K (1 mg ml^-1^) (HiMedia) or heat (50 °C for 1 hour). The treated cell-free supernatants were tested for their influence on biofilm formation or added into the wells of CR agar plated with a lawn of UPEC to determine its activity on matrix production.

### Biofilm assays and fluorescence microscopy

The microtiter plate assay in 96 wells was used to quantify the formation of partially submerged biofilm. Briefly, around 10^7^ mL^-1^ cells of UPEC were dispensed from overnight grown culture to microtiter wells containing YESCA broth and incubated for 24h at 37 °C in static condition. After rinsing to remove planktonic cells, the biofilm was stained by crystal violet (CV) and de-stained with 70% ethanol to quantify the biomass at 595 nm in a plate reader (Tecan Sunrise).

Dispersal studies of preformed biofilm was done according to Prasad *et al*., (2017), where the above-said procedure was followed to form biofilm on the surface of microtiter wells for 24 hours. Later, the media was decanted, the wells were rinsed thrice with PBS and 250 μl of the sterile supernatant from the isolates was added and incubated for 1 hour at 37 °C. For the control wells, 250 μl of the sterile PBS was added. Later, the residual biomass was stained with CV followed by de-staining and the absorbance were read at 600 nm to quantify the biomass.

Fluorescence imaging of biofilm was done according to Miryala *et al*. (2019). Nucleic acid stain, SYTO9 was used to stain the biofilm cells which was observed under fluorescence microscope (Nikon Eclipse Ni-U). Twenty randomly taken images were processed for auto-thresholding technique and the intensity (as proxy for biomass) and area coverage (substratum coverage) was measured in the ImageJ software (https://imagej.nih.gov/ij/index.html).

### Identification of the bacterial isolates

The bacterial isolates were identified by sequencing the 16S rRNA genes. Colony PCR was performed with 27f (GAGAGTTTGATCCTGGCTCAG) and 1541r (AAGGAGGTGATCCAGCCGC) universal primers. Amplicons were purified by standard procedures and sequencing was done by Eurofins India. Phylogenetic analysis was done using neighbor-joining method in the MEGA software version 7 (Kumar *et al*., 2016). Sequence-matched results were submitted to the GenBank and the bacterial isolates were submitted to National Centre for Microbial Resource, National Centre for Cell Science, Government of India (Table S1).

### Competition experiments

Competition experiments were performed in both planktonic and biofilm growth between the isolates and UPEC. The initial inoculum was 10^7^ CFU mL^-1^ for the monoculture or coculture experiments and it was performed in YESCA medium. For coculture experiments, the antibiotic sensitivity profile of selected isolates was tested and contrasting antibiotics were chosen for plating. UPEC was sensitive to ampicillin and the isolates B1, P1, and Z1 were resistant to ampicillin which was used for enumeration and calculation of fitness values. The isolates A1, C1 and T1 were transformed with pUltra plasmid (Mavridou *et al*., 2016) having gentamycin cassette and UPEC was transformed with pKD46 plasmid (Datsenko, and Wanner, 2000) having ampicillin cassette. Both UPEC and the bacterial isolate were mixed in 1:1 ratio in a centrifuge tube and along with the medium, dispensed in 24 well microtiter plate and, incubated at 25 ^o^C for 24h. After incubation, fitness was calculated for planktonic growth by plating in corresponding antibiotic containing media plates. For biofilm growth, the wells were rinsed thrice with PBS and scraped with a sterile rubber policeman to remove the biofilm cells that was plated on selective antibiotic plates for enumeration. Fitness was calculated as Malthusian parameter *M* = ln(*N*_1_/*N*_0_), where *N*_0_ is the initial cell number at 0 hour and *N*_1_ is the final cell number at 24 hours of incubation (Lenski *et al*., 1991). The absolute fitness was calculated for both monoculture and coculture between UPEC and the isolate. The relative fitness between monoculture and coculture and, also between the UPEC and the isolate in coculture experiments, were calculated by dividing monoculture/coculture and UPEC/isolate respectively.

The plasmids, pFPV expressing either GFP or cherry red (Valdivia and Falkow, 1996, Drecktrah *et al*., 2008) were transformed by electroporation into the isolate B1 and UPEC. Competition experiment was performed by inoculating the UPEC and the isolate in 1:1 (10^7^ CFU mL^-1^) ratio in a petri dish containing a glass slide with YESCA broth and incubated at 25 ^o^C for 24h. After incubation, glass slide was taken out, rinsed with PBS, dried and, observed under the fluorescent microscope (Nikon Eclipse Ni–U).

## Acknowledgements

CSS thanks financial support provided by Science and Engineering Research Board, Dept. of Science and Technology, Govt. of India (CRG/2018/001827) to conduct this study. We thank SASTRA Deemed University for the infrastructure, and SM and SC thanks the University for providing teaching assistantship (TA). We also thank DST-FIST funded fluorescence microscopy facility (SR/FST/ETI-331/2013).

## Authors’ contribution

SM and CSS conceptualized the study; SM and SC performed the experiments; SM and CSS analyzed the data and wrote the paper. All authors read and approved the final manuscript.

## Legends of Supplementary Materials

**Table S1**. Identity of the isolates with their corresponding NCBI and NCMR accession numbers

**Figure S1:** Influence of cell-free supernatant from colony of the isolates that were in contact with the UPEC colony on the (a) planktonic growth and (b) biofilm formation of UPEC.

**Figure S2:** Influence of different concentration of the Competition Sensing Independent Supernatant on UPEC biofilm formation

**Figure S3.** Influence of Competition Sensing Independent Supernatant (CSIS) of the isolates on UPEC (a) planktonic growth, (b) adhesion, (c) partially submerged biofilm (PSM) formation and, (d) dispersal of UPEC biofilm.

**Figure S4:** Quantified data of the fluorescence microscopic biofilm images (representative images shown in Figure 2e) of UPEC

**Figure S5**. Influence of physicochemical factors on the cell-free extract. (a) Representative image showing the color of the UPEC colony lawn in CR media indicative of matrix production. *n* = 2. (b) Planktonic growth and, (c) Biofilm formation of UPEC in presence of cell-free extract treated with BME, proteinase K, trypsin and, heat at 50 °C. *n* = 5

**Figure S6.** Absolute fitness values of the isolate and UPEC in monoculture (mono) and coculture (co) in both planktonic growth and biofilm growth. (a) Isolate A1, (b) Isolate B1, (c) Isolate C1, (d) Isolate P1, (e) Isolate T1, and (f) Isolate Z1

**Figure S7.** Absolute values of biomass and substratum coverage of the monoculture and coculture biofilm of UPEC and *S. enterica* B1, quantified from the fluorescence images. (a) Biomass and (b) Substratum coverage. Black boxes represent total biomass and coverage, red boxes are the UPEC and green represents the *S. enterica* B1.

