## Supplementary figures and images for "Competitive exclusion of uropathogenic *E. coli* biofilm by *Salmonella* through matrix inhibition"

### Figure S1

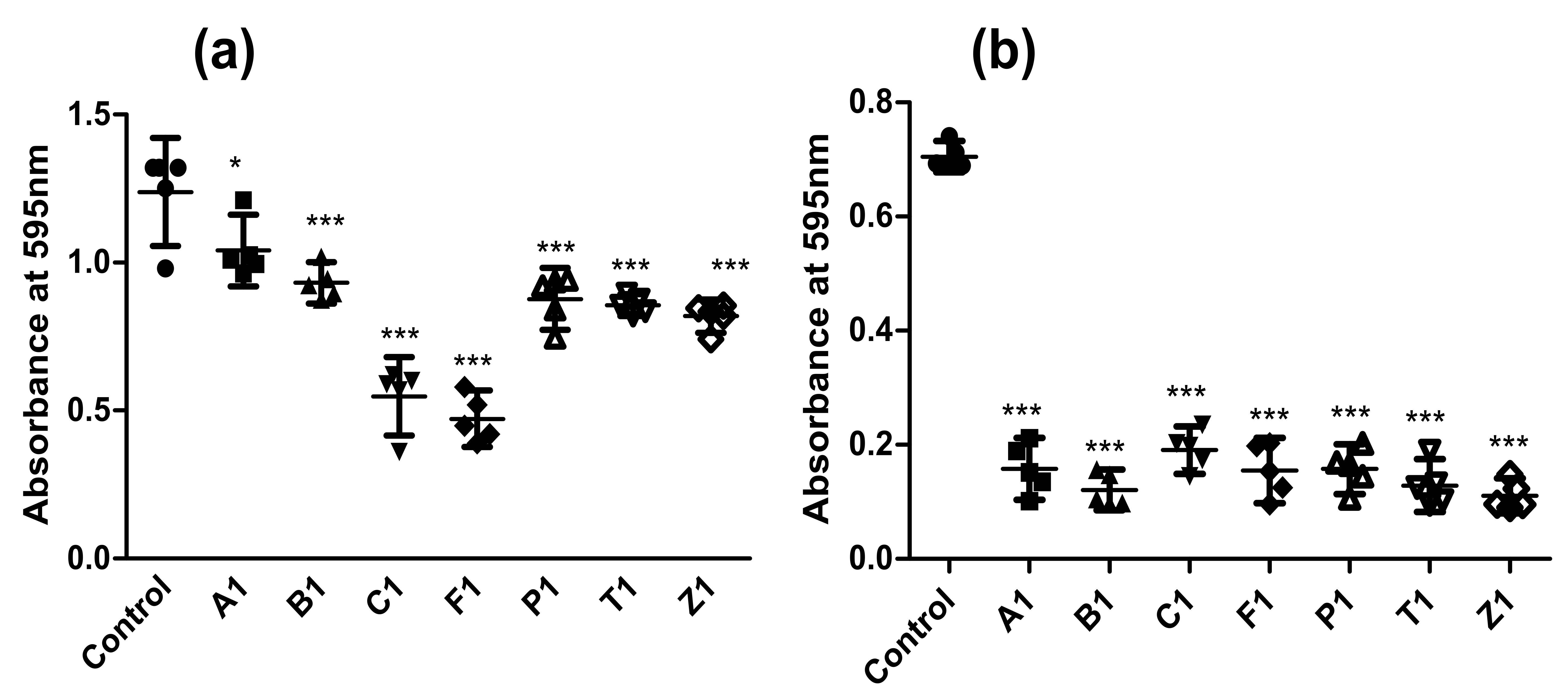

### Figure S2

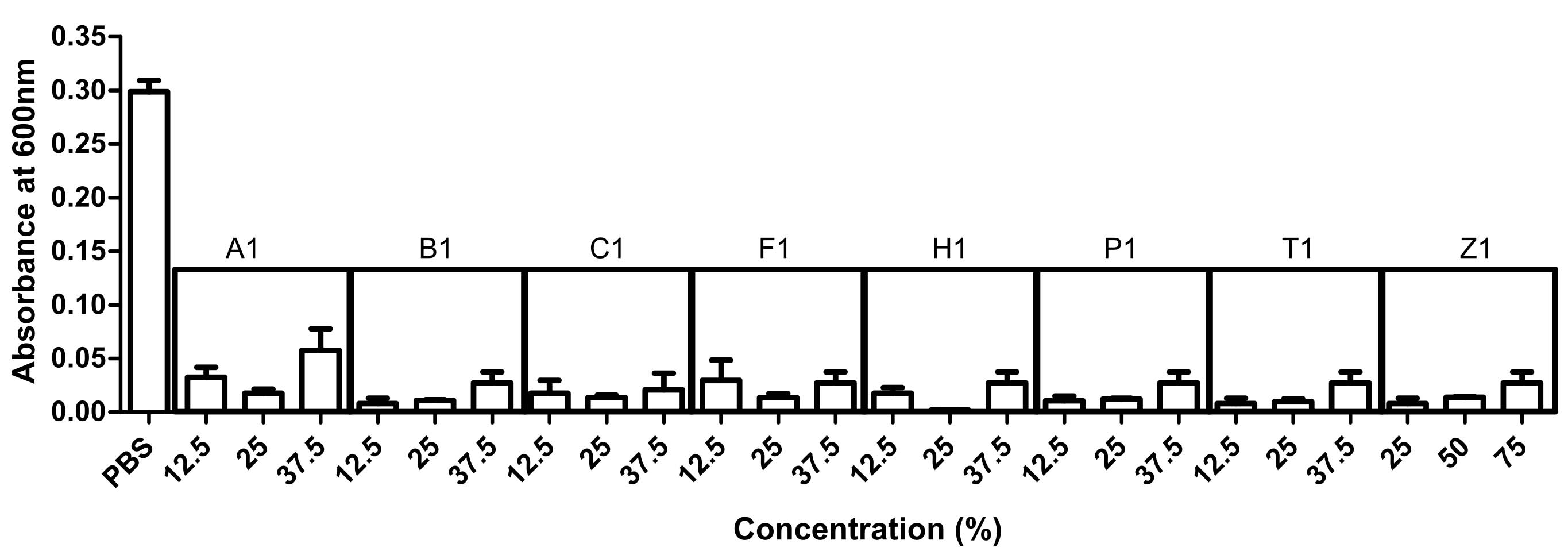

### Figure S3

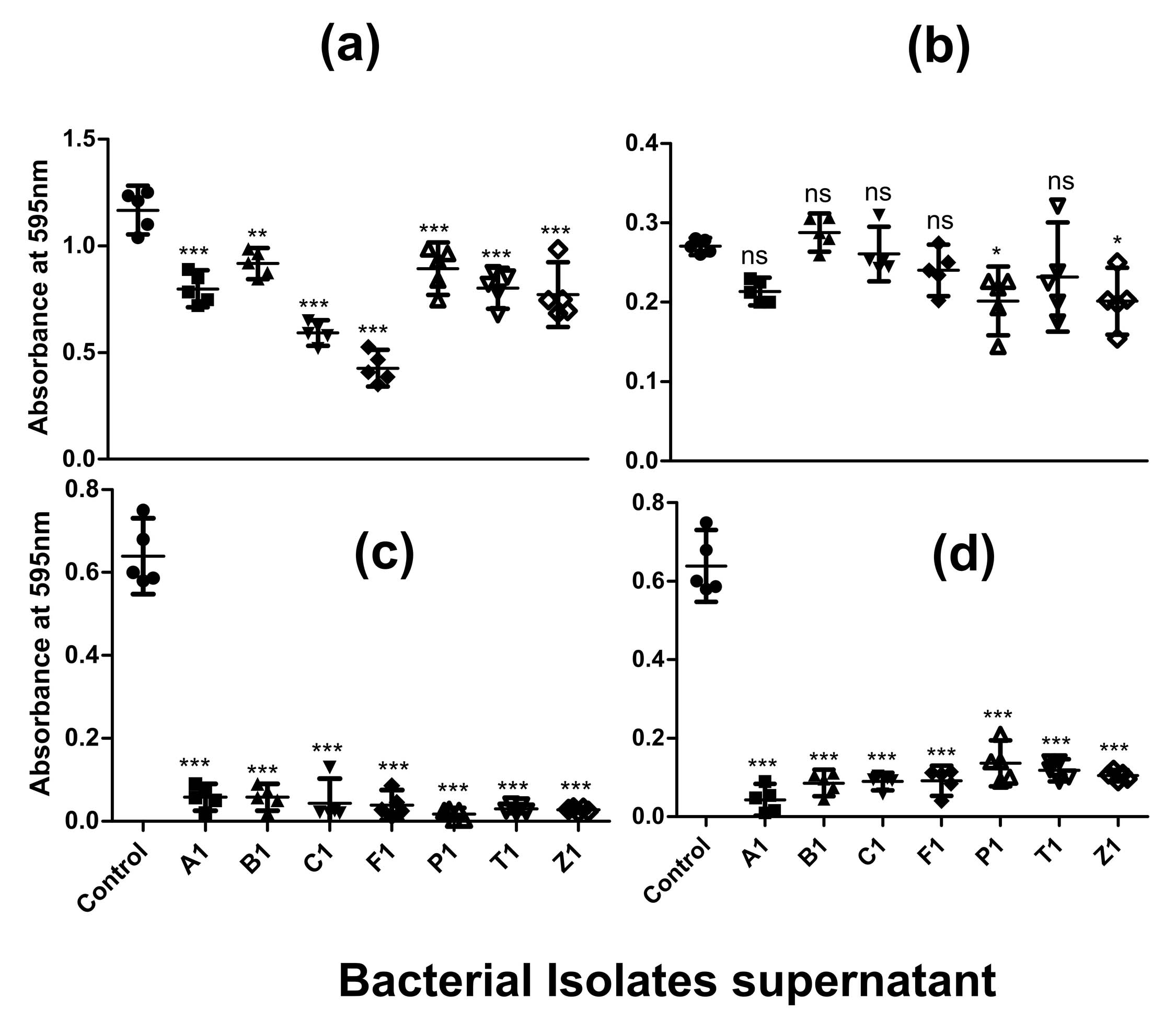

### Figure S4

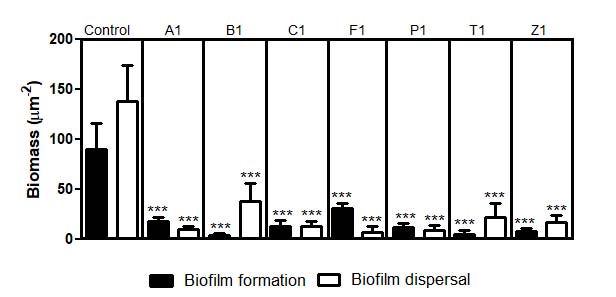

### Figure S5

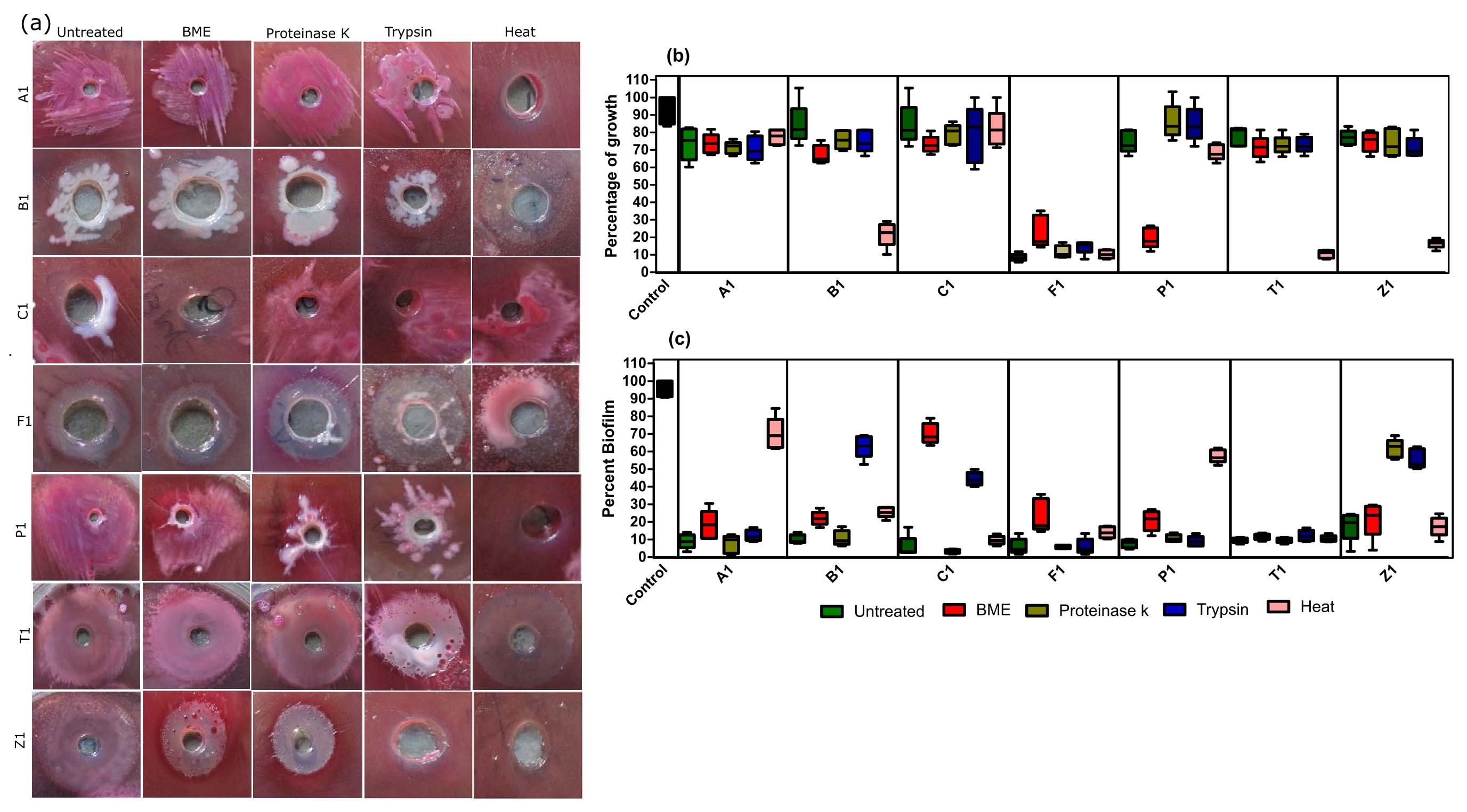

### Figure S6

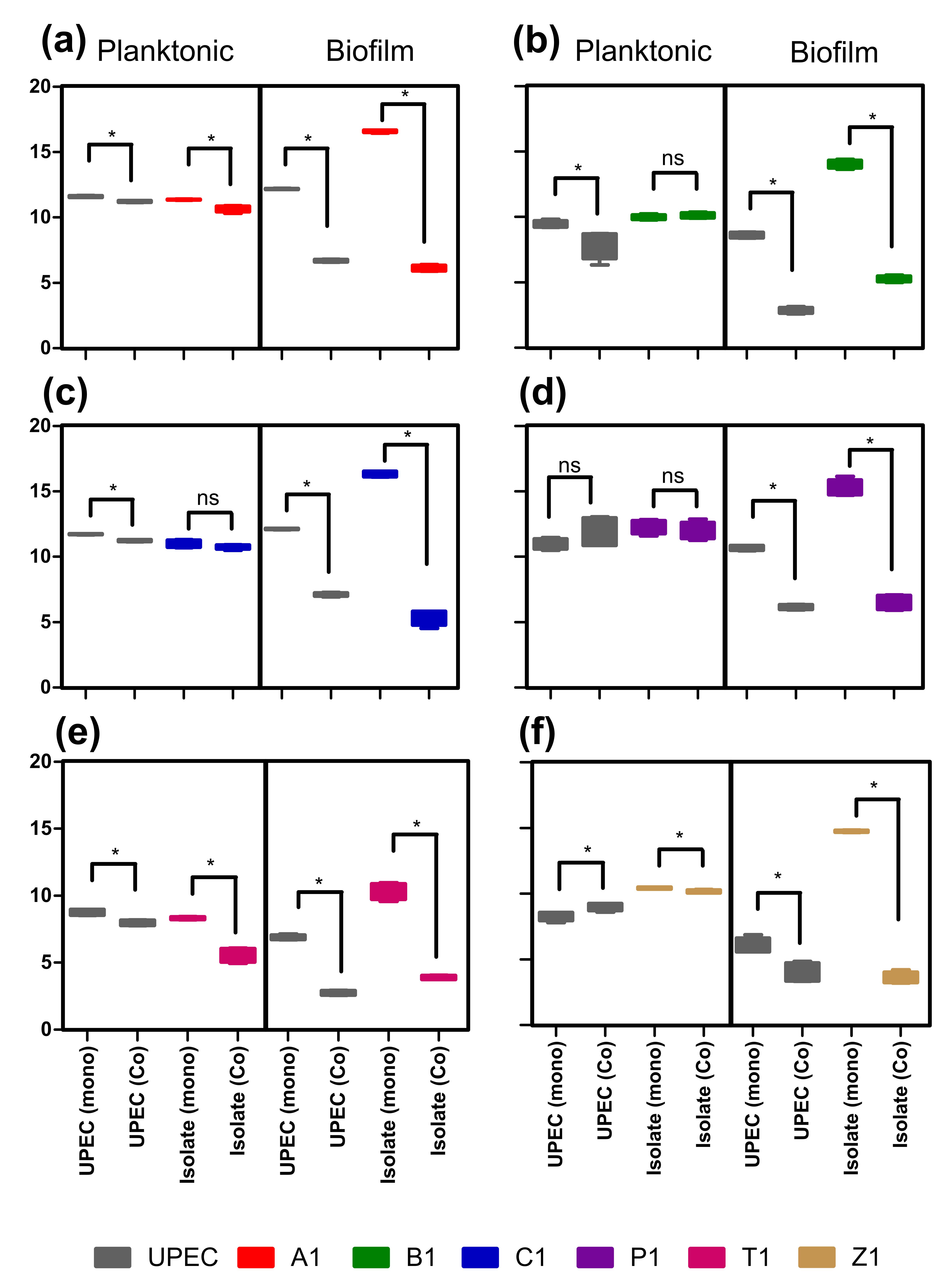

### Figure S7

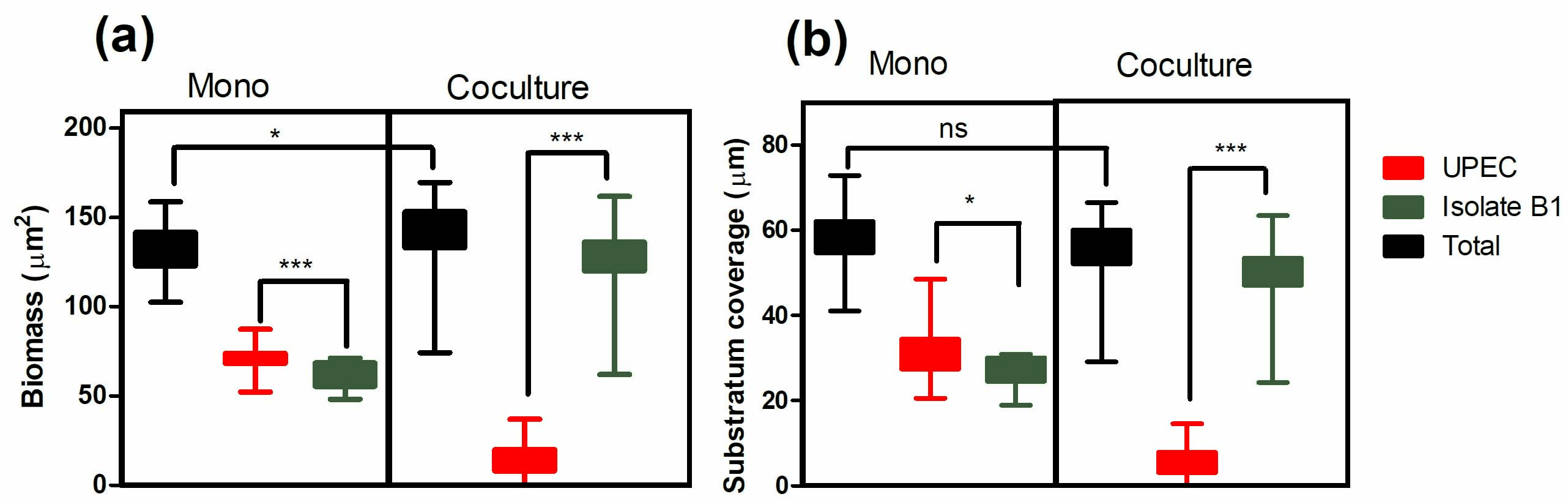
