## Supplementary material for "Competitive exclusion of uropathogenic *E. coli* biofilm by *Salmonella* through matrix inhibition": Table S1

**Table S1**. Identity of the isolates with their corresponding NCBI and NCMR accession numbers

| **Isolate name** | **Identity** | **NCBI***  **accession number** | **NCMR** accession number** |
| --- | --- | --- | --- |
| A1 | *Escherichia fergusonii* | MG190339 | MCC 4245 |
| B1 | *Salmonella enterica* | MG062784 | Submitted |
| C1 | *Escherichia* sp. | MG182420 | Submitted |
| F1 | *Escherichia fergusonii* | MN180846 | Submitted |
| P1 | *Shigella flexneri* | MG062784 | MCC 4276 |
| T1 | *Shigella flexneri* | MK817649 | MCC 4244 |
| Z1 | *Enterobacter cloacae* | MN186811 | MCC 4253 |

* National Center for Biotechnology Information, US

** National Centre of Microbial Resource, India
